# Familial Risk for Dementia, Cognitive Performance, and the Human Cerebello-Hippocampal Circuit: A Study in the European Prevention of Alzheimer’s Disease cohort

**DOI:** 10.1101/2025.09.21.677596

**Authors:** Tracey H. Hicks, Jessica A. Bernard

**Affiliations:** Department of Psychological and Brain Sciences, Texas A&M University, College Station, Texas, United States of America; Texas A&M Institute for Neuroscience, Texas A&M University, College Station, Texas, United States of America

**Keywords:** functional connectivity, cerebellum, hippocampus, cognition, dementia risk, sex differences

## Abstract

The cerebello-hippocampal (CB-HP) circuit is increasingly implicated in episodic and spatial memory, yet its role in normal aging, dementia risk, and sex differences remains unclear. Structure and function in both the hippocampus and cerebellum have been linked to mild cognitive impairment, Alzheimer’s Disease, and cognitive decline in healthy older adults. Literature on the CB-HP circuit is largely limited to animal studies and small samples of young adults in the context of visuospatial abilities, underscoring the need for large, diverse aging cohorts to establish CB-HP connectivity as an early, translatable marker of neurodegeneration. Here, we investigate the CB-HP circuit in the context of aging, dementia risk, behavior, and sex differences in healthy older adults to advance understanding of cognitive decline in the aging population. We explored this relationship in 857 healthy adults with and without a family history of dementia (aged 50-88, 59% female, 71% family history positive) using resting-state functional connectivity MRI (rsMRI) and behavioral assessments. We hypothesized that CB-HP functional connectivity (FC) would be lower with increased age, relate positively to cognitive performance, be lower in females positive for family history of dementia, and be lower in mild cognitive impairment (MCI)-risk than cognitively normal (CN) participants with sex differences therein. We observed selective links between CB-HP FC, cognition, and sex. MCI risk participants performed worse on spatial memory performances than CN, whereas CB-HP FC showed opposite performance slopes by risk status, suggesting the CB-HP circuit may function as a compensatory network for short-term recall. Sex differences were seen on cognitive tasks (delayed episodic memory and spatial tasks) and in CB-HP a better visuospatial index was linked to greater FC in females, while males displayed an inverse relationship. Behavioral differences by familial dementia history were shown, although CB-HP FC did not show effects here. Overall, CB-HP networks appear behaviorally silent in the aggregate but reveal risk- and sex-dependent relationships when memory demands and circuit nodes are considered.

## Introduction

Cognitive and motor abilities decline with healthy aging and are dramatically impacted by neurodegenerative diseases like Alzheimer’s disease (AD)^1–3^. Reliable and accurate identification of early pathology in aging is crucial for developing interventions for healthy decline and disease-related pathology. Structure and function in both the hippocampus and cerebellum have been linked to mild cognitive impairment^4,5^, AD^4,6–10^, and cognitive decline in healthy older adults (OA)^11–16^. Notably, studies have demonstrated synchronized neuronal oscillations, coactivation, and functional connectivity (FC) between these two regions^17–19^. Further support for direct communication between these regions are shown in both human and animal studies^20–25^ which has led to the exploration of the cerebello-hippocampal (CB-HP) circuit. The CB-HP circuit has been associated with lower FC with increased age^19^. Conversely, Paitel et al. linked broader cerebello-hippocampal FC (i.e., right and left hippocampus FC to right and left cerebellum FC; processed as 4 regions) to greater FC with increased age in healthy adults across middle and older age; however, they also showed bidirectional age effects with greater CB-HP FC in (apolipoprotein E)APOE ε4 non-carriers and lower FC in ε4 carriers or individuals with abnormal tau/amyloid ratios with increased age^18^. Given that cognitive and motor abilities deteriorate with age, the integrity of the CB-HP circuit may represent a promising biomarker for pathology in healthy aging or prodromal dementia risk^26^. Examining familial history of dementia alongside this circuit stands to provide distinct insights from genetic CB-HP associations. Extant literature on the CB-HP circuit is largely limited to animal studies^20–22^ and small samples of young adults in the context of visuospatial abilities^23–25^, underscoring the need for large, diverse aging cohorts to establish CB-HP connectivity as an early, translatable marker of neurodegeneration.

Specific types of cognitive tasks have been linked to cerebello-hippocampal coactivation^25^ and cognitive decline^1–3^, respectively. Human studies have demonstrated cerebello-hippocampal coactivation during tasks which require time and space sensitive prediction of movements with visuomotor integration^25^. Although these insights naturally apply to spatial and cognitive-motor functioning, they also implicate mechanisms of episodic memory^17^, such that autobiographical events are stored as temporally ordered sequences or conceptual “routes” through life, mirroring how we map physical places and paths (e.g., milestones)^27^. Both episodic memory and spatial abilities exhibit deficits in early-stage AD^28–30^, implying that the CB-HP circuit plays an integral role in certain cognitive processes that are impacted during neurodegeneration. In a broader examination of cerebello-hippocampal FC, Paitel and colleagues found greater FC between both the right hippocampus and left cerebellum as well as right hippocampus to right cerebellum were associated with better episodic memory performance in healthy aging adults^18^. Here, the goal is to determine the role of CB-HP interactions as they relate to specific cognitive tasks (e.g., spatial processing, episodic memory, and spatial memory) in a large sample of older adults (n=857). This will test the hypothesis that CB-HP interactions at rest are associated with specific cognitive processes.

Both a family history of dementia (FH+) and biological sex (i.e., female) have been identified as unique risk factors for AD^31–37^. There is growing evidence to support the theory that dementia pathology starts 10-20 years before the disease is diagnosed^36,38–41^. Cognitively unimpaired middle-aged adults who carry this family history already exhibit indicators of hallmarks seen later in clinical cohorts such as hippocampal atrophy^41–43^ and disrupted functional networks^44,45^. Numerous studies have shown FH+ is an independent risk factor for dementia with effects that are partly dissociable from genotype-based susceptibility^46–49^, implying that FH+ will provide a uniquely informative lens on preclinical markers. Because 71% of our cohort is FH+, the present study is well positioned to interrogate this factor in depth. Female sex represents an intersecting vulnerability: females have a higher lifetime prevalence of dementia^50,51^ and experience abrupt endocrine transitions that impact FC^52–54^. These converging lines of evidence position FH+ and female sex as critical modulators of brain aging and underscore the need to probe their impact on the CB-HP circuit for use as a sensitive barometer of prodromal neurodegeneration^19,26^. This study characterizes the CB-HP circuit in healthy older adults with and without risk factors for AD as it relates to cognitive performance with the intent to identify early indicators of cognitive deficits that translate to clinical use.

Global measures of cognitive function are frequently used to identify cognitive deficits and risk for developing dementia^55,56^. The Repeatable Battery for the Assessment of Neuropsychological Status (RBANS)^57^ gauges global cognitive function and was designed to detect very mild dementia^56^. The Total Scale Index (TIS) is derived from subtests in several cognitive domains that have been standardized to a normative sample and TIS scores have shown efficacy in discriminating between mild cognitive impairment (MCI) and cognitively normal (CN) adults, with receiver-operating curves indicating 84% sensitivity and 90% specificity^56^. These empirically derived groupings provide a framework for probing early neural and cognitive change. Integrating these RBANS-defined categories into our investigation of the CB-HP circuit and cognitive performance therefore represents a critical next step for investigating who is most vulnerable to dementia.

This study aimed to characterize the CB-HP circuit in the context of cognitive performance and dementia risk in older adults. CB-HP FC relationships with cognition were evaluated across domains of episodic memory, spatial memory, and spatial processing speed. We hypothesized that cognition would display positive relationships with CB-HP FC (i.e., better performance associated with greater FC). Sex differences were assessed across participants and within groups (FH+ and RBANS, respectively). We predicted FH+ females would show reduced CB-HP FC as compared to FH+ males. In our exploratory analyses of RBANS CN and MCI risk groups, we expected that MCI risk groups would exhibit lower CB-HP FC as compared to CN and sex differences therein. Finally, we hypothesized that cognitive performance would be lower in participants with dementia risk (i.e., females, FH+, and the RBANS MCI risk group). Within those results, we expected measures of spatial memory task performance to be lower in those at risk for dementia.

## Methods

### 1.1. Study sample

Participants were from the European Prevention of Alzheimer’s Disease (EPAD) cohort^58^. There were criteria set by a balancing committee (BC) to estimate the probability of a participant developing dementia and participants were recruited with the design of developing a successful dataset for disease modeling. That is, EPAD includes healthy participants with no risk factors for developing dementia as well as participants with a variety of risk factors. To be eligible they must be aged 50 years or older, able to read, write, have a minimum of 5 years of formal education, and have a study partner (collateral) for the screening visit. For all participants exclusion criteria include: 1) diagnosis of dementia at baseline; 2) a Clinical Dementia Rating (CDR) score ≥ 1 at baseline; 3) Presenlin (PSEN) PSEN1, PSEN2, or APP mutation carrier; 4) Presence of any neurological, psychiatric or medical conditions associated with a long-term risk of significant cognitive impairment or dementia including but not limited to pre-manifest Huntington’s disease, multiple sclerosis, Parkinson’s disease, Down syndrome, active alcohol/drug abuse; or major psychiatric disorders including current major depressive disorder, schizophrenia, schizoaffective or bipolar disorder; 5) Any cancer or history of cancer in the preceding 5 years; 6) Any current medical conditions that are clinically significant and might make the subject’s participation in an investigational trial unsafe; 7) Any contraindications for MRI/positron emission tomography (PET) scan; 8) Any contraindications for lumbar puncture at visit 1 (including refusal of lumbar puncture procedure to collect sample); 9) Any evidence of intracranial pathology which, in the opinion of the investigator, may affect cognition; 10) Participation in a clinical trial of an investigational product (CTIMP) in the last 30 days; 11) Diminished decision-making capacity/not capable of consenting at Visit 1; and 12) Unable to comply with protocol requirements in the opinion of the investigator. The current analyses were utilized the EPADLCS-v.IMI dataset, released in October 2020 and accessed from the Workbench of the Alzheimer’s Disease Data Initiative (ADDI) Portal. EPAD LCS is registered at www.clinicaltrials.gov Identifier: NCT02804789. The dataset includes data from all participants who consented to join the study. Data used in preparation of this article were obtained from the EPAD LCS data set V.IMI, doi:10.34688/epadlcs_v.imi_20.10.30. This study evaluated the baseline (V1) data which has 1906 participants; however, this study focused on participants with preprocessed neuroimaging data (*n*=857).

All participants underwent a baseline visit which included a battery of cognitive tasks and a magnetic resonance imaging (MRI) session amongst other variables that were collected. Cognitive data is primarily derived from V1; however, Visit 2 (V2) cognitive data were used for the Virtual Supermarket Task and 4 Mountains test if data from V1 was missing. Here, we focused on rsfMRI, cognitive performance, and relationships with sex and family history of dementia.

Demographic details are presented in **Table 1**. Family history of dementia was examined via dummy variable, *parent history of dementia*, which broke down participants into two groups, positive for one or both parents having a history of dementia (FH+) and negative for any parent history of dementia (FH-). These groups were then matched by age and sex using the ‘matchit’ package in R^59^. Ethnicity was included in **Table 1** for generalizability purposes and we recognize that racial and ethnic demographics are social and political categories given meaning by social, historical, and political forces^60^. Race and ethnicity will not be evaluated separately in this study, as information about socio-economic status is better indicated for causal inferences in neuroimaging^60^. It is also notable that demographics in European countries have greater focus on country membership as compared to the United States’ practice of examining ancestral origin to distinguish demographics. Eurostat reports demographics at the European Union level by year with respect to numbers of residents and immigration levels; however, this information cannot be directly applied to determine whether the EPAD dataset accurately represents the ethnic distribution obtained.

**Table 1.** Demographic means for the sample and sub-samples are listed below and ethnicity is reported in percentages.

|  | Demographics |  |
| --- | --- | --- |
| **Full sample** | Age (years) | 64.95±7.28 |
|  | Education (years) | 14.50±3.88 |
|  | Sex F(M) | 508(349) |
|  | African (%) | 0.2% |
|  | Asian (%) | 0.6% |
|  | Caucasian (%) | 77.1% |
|  | Not reported (%) | 21.4% |
| **Family History of Dementia** | **FH+** | **FH-** |
| Number of Total Participants | 605 | 252 |
| Number of Age/Sex Matched Participants | 239 | 239 |
| Age (years) | 66.5±7 | 66.5±7 |
| Education (years) | 14.5±4 | 14.9±3.8 |
| Sex F(M) | 132(107) | 132(107) |
| **RBANS Group** | **MCI risk** | **CN** |
| Number of Participants | 104 | 748 |
| Age (years) | 68±7.9 | 64.5±7.1 |
| Education (years) | 11.5±3.8 | 14.9±3.7 |
| Sex F(M) | 55(49) | 451(297) |
|  | African (%) | 0.2% |
|  | Asian (%) | 0.6% |
|  | Caucasian (%) | 77.1% |
|  | Not reported (%) | 21.4% |

*Note.* Family history group demographics are listed for the age and sex-matched groups that were created (i.e., the data that FH+/- analyses were conducted with). The number of total FH+/- participants is also listed for context.

All study procedures conducted by these authors were approved by the Institutional Review Board at Texas A&M University. EPAD participants consented for their data to be used in future research projects and data from the EPAD LCS is open access. All data for this project were stored on secure databases. Specifically, ADDI Workspaces and Texas Virtual Data Library (ViDaL; see Equipment for additional details).

### 1.2. Cognitive Testing

Participants completed a battery of cognitive tasks which quantify processing speed, visuospatial abilities, and episodic memory.

#### 1.2.1. Repeatable Battery for the Assessment of Neuropsychological Status

RBANS is a standardized instrument designed to assess cognitive functioning across five domains: immediate memory, visuospatial/constructional, language, attention, and delayed memory and also yields a Total Scale Score, which provides an estimate of overall cognitive functioning^57^. The RBANS has demonstrated good sensitivity and specificity for detecting cognitive impairment in older adults and is frequently used in both clinical and research settings to evaluate cognitive decline associated with aging and neurodegenerative conditions^56^. For this study we used the RBANS total scale index, individual index scores (immediate memory, delayed memory, and visuospatial constructional). The total index score was used in our study to evaluate for global neuropsychological status^61^. Cutoff scores derived from Karantzoulis and colleagues (2013), were used to proxy a healthy cognitively normal (CN) profile (RBANS total scale index greater than or equal to 85) and a mild cognitive impairment (MCI) risk profile (RBANS total scale index less than 85)^56^. We also evaluated individual measures: Immediate Story Memory, Delayed Story Memory, and coding. Cognitive domain specific index scores and individual tests were evaluated to assess nuanced spatial abilities and episodic memory as it relates to a family history of dementia and the CB-HP circuit.

#### 1.2.2. 4 Mountains Test

The 4 Mountains Test^62^ is a brief, non-verbal test of allocentric spatial memory designed to detect early signs of Alzheimer’s disease. Participants are asked to identify the same mountainous landscape from a different viewpoint after a short delay, challenging their ability to form and recall spatial representations independent of perspective. Our study examined the percent of correct responses on this test as the outcome, with a higher percentage indicating better performance. The test has demonstrated high sensitivity in distinguishing individuals with pre-dementia Alzheimer’s disease from healthy controls^62^, making it a promising tool for early detection.

#### 1.2.3. Virtual Supermarket Task

Spatial orientation was assessed using a validated virtual reality paradigm known as the Virtual Supermarket Task^63^, which provides an ecologically relevant and non-verbal measure of egocentric spatial memory. Participants viewed short video clips (from a first-person perspective) simulating travel through a supermarket while pushing a shopping cart, involving a series of 90° turns to various locations. After each trial, participants were asked to indicate the direction of the original starting point. The environment lacked salient landmarks, requiring incidental encoding of the route and continual spatial updating. The task consisted of 14 trials (two blocks of 7), each standardized for route complexity and duration. A brief practice trial was administered prior to testing, and directional responses were recorded on two axes (left/right and front/behind), with overall accuracy calculated based on correct responses across both dimensions. Here, we used the percent of correct responses to assess performance on this task and converted them to z-scores to standardize performance within the sample. A higher z-score on this task indicates better performance.

### 1.3. Imaging acquisition and Processing

Participants underwent structural and resting-state MRI using Siemens and Philips scanners across 15 research sites. Lorenzini and colleagues have provided detailed documentation of the protocols, dataset, and processing workflow^64^. Each participant had 1 rsfMRI scan. In the EPAD LCS protocol (v5.0), MRI acquisition is split into core sequences obtained for all participants and advanced sequences collected only at sites with the requisite technology and experience; rsfMRI is part of the advanced set. Core imaging comprised 3D T1-weighted and 3D FLAIR structural scans acquired at all sites; advanced imaging included rs-fMRI. For rs-fMRI, vendor-matched parameters were harmonized: Siemens EPI with voxel size 3.3×3.3×3.3 mm³, TR = 2020 ms, TE = 30 ms, 204 volumes, A→P phase-encode, AT = 6:52; Philips EPI with voxel size 3.3×3.3×3.3 mm³, TR = 1640 ms, TE = 30 ms, 202 volumes, A→P phase-encode, AT = 5:35; rs-fMRI was not acquired at the GE site. Multiband/SMS factors were not specified in the EPAD acquisition table. Structural imaging used vendor-specific 3D T1w scans with approximately 1.0–1.2 mm isotropic voxels (e.g., Siemens: 1.20×1.05×1.05 mm³, TR = 2300 ms, TE = 2.95 ms), and 3D FLAIR at 1.0 mm isotropic; additional core 2D T2w/T2* sequences were acquired for radiological assessment. rsfMRI underwent reversed phase-encode distortion correction (FSL topup), motion correction, and registration to T1w; processing was performed in ExploreASL with SPM12 and FSL.

The anatomical and functional images were preprocessed using fMRIPrep version 21.0.1^65^, which includes automated procedures to align the functional volume with the anatomical image, correct for motion, correct field map distortions, segment the anatomical image into distinct tissues (e.g., gray matter, white matter, cerebrospinal fluid), remove the skull from the anatomical image, normalize the data to a common space, align motion-corrected functional volumes with the normalized anatomical image, and apply spatial smoothing. Esteban and colleagues have published additional details on the fmriprep preprocessing pipeline^65^.

### 1.4. Neuroimaging analysis

This section utilized standardized text to ensure consistency and reproducibility in reporting the research protocol across work completed by our group^19,66,67^. Following preprocessing with fMRIPrep, the subsequent analyses were conducted using the CONN toolbox (version 21a)^68^. This involved additional processing to eliminate noise and artifacts and enhance data quality. Denoising in CONN typically comprises several stages, such as removing motion signals and regressing out confounding signals (e.g., signals from ventricles, white matter, and global signals). Motion information from fMRIPrep was utilized in CONN for this purpose. A 0.008-0.099 Hz bandpass filter was applied to eliminate high-frequency noise. The denoising step is crucial for enhancing the quality of FC data by minimizing artifacts and enhancing the ability to detect genuine FC patterns in the brain.

Resting-state FC analyses focused on regions of interest (ROIs) in both the hippocampus and the cerebellum (**Figure 1**). The hippocampal seed regions were loosely based on coordinates derived from Grady’s meta-analysis on hippocampal function^69^. These coordinates helped determine anatomical hippocampal boundaries and the original coordinates were shifted from the original values to accommodate a new center for 5mm spherical seeds in MNI space. That is, certain coordinates from Grady^69^ fell on the outer edges of the hippocampus and to cover 3-dimensional space within the boundaries of the hippocampus, we had to shift certain coordinates to account for a 5mm spherical seed. Cerebellar seeds were determined by mapping dorsal, ventral, rostral, caudal, medial, and lateral boundaries of the right cerebellum in MNI space. Please refer to **Supplementary Table 1** for coordinates for all 52 our seeds. Following this step, we used the oro.nifti and spatstat packages in R Studio to create a non-overlapping grid in 3 dimensional space of 5mm spherical seeds covering the right cerebellum^70,71^. As our aim for this study was to cover the entirety of the right cerebellum and left hippocampus, seeds were created to cover the 3-dimensional anatomical regions in MNI space. For ease of interpretation, we used the automated anatomical atlas (AAL) in the label4MRI package in R to approximate anatomical regions for each seed^72^ which is specified in **Supplementary Table 1**. Our seeds are visualized in a simpler, easier to view, rendering in **Figure 1**, and we have created a more in-depth representation in **Supplementary Figure 1**. These seeds have also been used in a study of the CB-HP circuit in middle-aged and older adults^19^; our intention was to run parallel analyses with the same seeds across large, healthy adult cohorts to characterize the CB-HP circuit across the lifespan. The investigation was limited to the right cerebellar hemisphere and left hippocampus, to mitigate multiple comparisons and ensure examination of cross lateralized connections.

**Figure 1.**
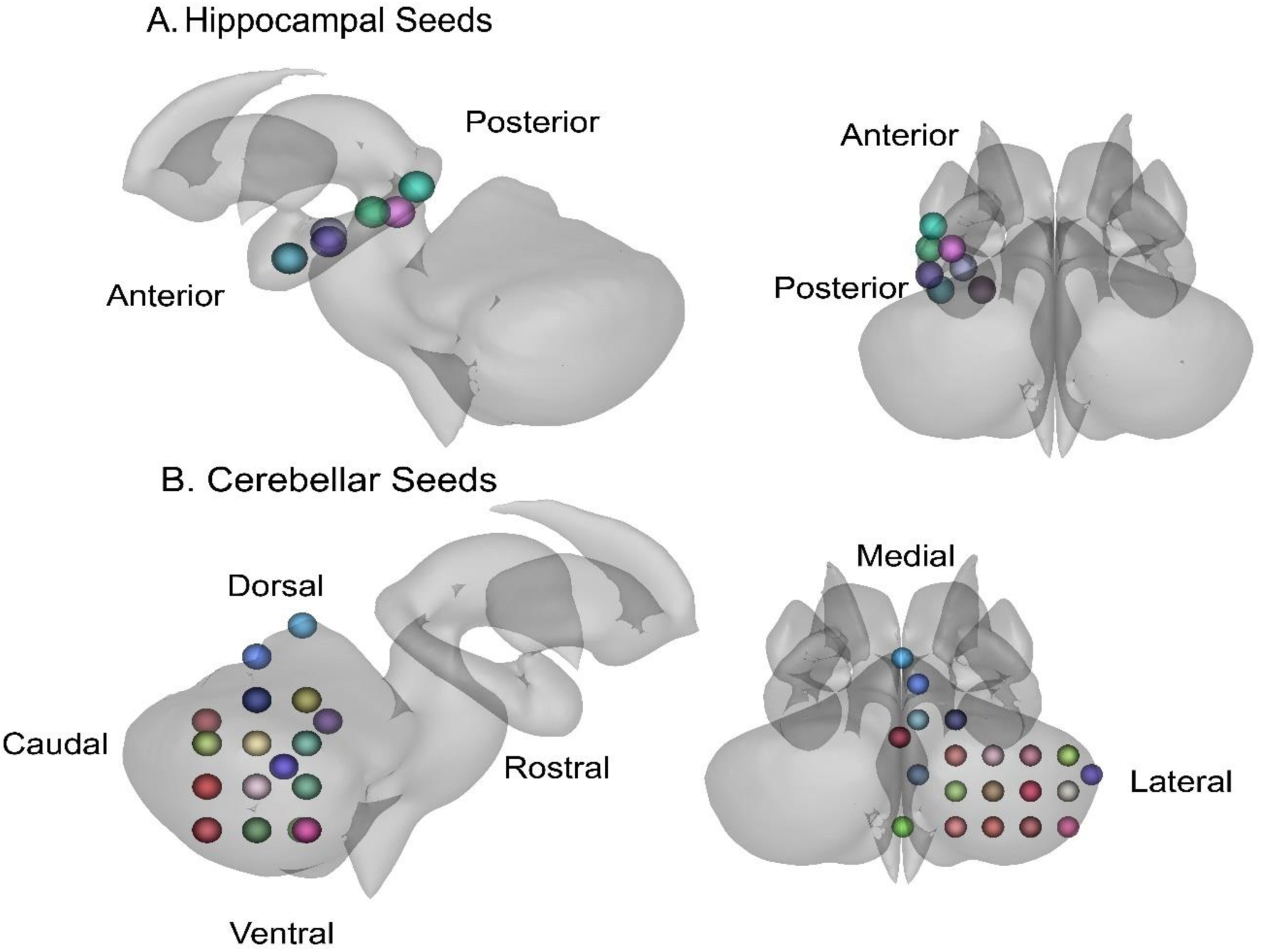
Seed regions used in this study. Seed colors are helpful for differentiating seeds, but do not hold additional meaning. **A**. Hippocampal seeds are displayed with directional indicators of the anterior and posterior long-axis. **B**. Cerebellar seeds are displayed with directional indicators for the right cerebellar hemisphere. The caudal region of the cerebellum is better visualized in a more detailed representation of the ROI seeds in **Supplementary Figure 1**.

CB-HP FC was examined via region of interest (ROI)-to-ROI correlations using CONN toolbox at the group level. We used an initial voxel threshold at p<.001 along with a cluster threshold set at p < .05, with a false discovery rate (FDR) correction. To determine effect size specifically with Cohen’s d, we utilized the equation ((2* t-value)/√degrees of freedom)^73^ in R Studio. To reduce inter-site variability in rsfMRI FC measures, the ComBat harmonization was implemented using the neuroCombat function from the neuroCombat R package^74^, which applies an empirical Bayes framework to adjust for scanner/site effects while preserving variance related to biological covariates of interest^75^. ComBat harmonization was conducted with scanner site as the batch variable and covariates included: age, education level, family history of dementia, and each cognitive measure. To conduct harmonization, ComBat requires all cells to be filled. 15 ROI-to-ROI pairs were missing values for the ROI-to-ROI correlations (listed in **Supplementary Table 2**). We excluded these correlations from our analyses. All of the following analyses will refer to ROI-to-ROI correlations with the implication that these relationships were evaluated with ComBat harmonization. FDR correction was also applied to all the following analyses.

Due to our robust multiple comparisons corrections, we also conducted exploratory analyses with a subset of ROI-to-ROI pairs. Specifically, we examined 22 ROIs defined as Crus I, Crus II, and the hippocampus in the AAL atlas^72^. These cerebellar regions were chosen for the subset analyses due to their associations with cognitive performance in the literature^21,24,25,76,77^. All the following FC analyses were run with the full set of ROI-to-ROI pairs first and then the subset of pairs. In the full set of ROI pairs, FDR was correcting for 1,320 variables and in the subset FDR corrected for 220 variables.

To determine cognitive performance relationships with CB-HP FC, the variables story memory immediate recall, story memory delayed recall, coding, immediate memory index, delayed memory index, visuospatial/constructional index, 4 Mountains, and Virtual Supermarket Task were separately examined with CB-HP FC via linear regressions. To determine whether education had an impact on these relationships, follow-up analyses on the variables story memory immediate recall, story memory delayed recall, coding, 4 Mountains, and Virtual Supermarket Task (which did not have education level controlled for in normative data) were separately correlated with CB-HP FC while controlling for education level.

Relationships between age, education level, and CB-HP FC were evaluated via linear regressions in the whole sample with CB-HP as the outcome. We assessed for sex differences within the parent history of dementia groups via linear regressions with the FH groups and sex variables as an interaction (predictor) and CB-HP as the outcome in a subset of age and sex matched participants. Age was matched within 1 year (e.g., participant age 64.4 would be matched to a participant between 63.4 and 65.4). 379 participants were excluded from the analyses to produce this age and sex matched sub-sample. To determine interactions between FH group and cognitive performance relationships with CB-HP FC, the variables story memory immediate recall, story memory delayed recall, coding, immediate memory index, delayed memory index, visuospatial/constructional index, 4 Mountains, and Virtual Supermarket Task were separately evaluated via linear regressions with CB-HP FC as the outcome and the interaction between cognitive performance and FH group as the predictor in an age and sex matched sample.

To explore another facet of preclinical cognitive decline in our healthy population, we examined RBANS Total Index Scores (TIS) with a cutoff score to look at MCI risk (<85) versus CN (85+) participants (previously explained in greater detail in Methods section 1.2.1) using linear regressions with CB-HP FC as the outcome and both main effects and interactions between RBANS group and cognition (on all tasks separately) as the predictors; these analyses were replicated to control for education level for tasks without normative data correction. FDR correction here accounted for 1,320 comparisons in the full set of ROIs and 220 comparisons in the subset of ROIs in each unique regression.

### 1.5. Statistical analyses of Demographics and Behavior

For statistical analyses of the age, education, sex, cognitive performance, and family history results, we used R (v2024.04.2.764, R Core Team, 2021). FDR correction was also conducted in R across all statistical analyses to account for multiple comparisons.

We first investigated interactions and main effects of the predictor parent history of dementia (i.e., one or both parents with a dementia diagnosis (FH+) versus no parent with a dementia diagnosis(FH-)) and binary sex, with the outcome cognitive performance (i.e., story memory immediate recall, story memory delayed recall, coding, immediate memory index, delayed memory index, and visuospatial/constructional index, 4 Mountains, and Virtual Supermarket Task) via separate linear regressions in an age and sex matched subset of participants; education level was controlled for in cognitive tests without normative data correction (story memory immediate recall, story memory delayed recall, coding, 4 Mountains, and Virtual Supermarket Task). Sex differences as they relate to cognitive performance (i.e., story memory immediate recall, story memory delayed recall, coding, immediate memory index, delayed memory index, and visuospatial/constructional index, 4 Mountains, and Virtual Supermarket Task) were assessed via linear regressions with sex as a binary predictor, controlling for age and education in tasks without normative data correction.

RBANS total index scores were converted to a dummy variable to explore MCI risk (<85) and CN (85+) participants’ differences and binary sex differences within cognitive performance on specific subtests and indices of the RBANS (story memory immediate recall, story memory delayed recall, coding, Immediate memory Index, delayed memory index, and visuospatial/constructional index) as well as egocentric and allocentric measures of spatial memory (4 Mountains and Virtual Supermarket Task) via interactions and main effects in linear regressions with RBANS group and sex as binary predictors. We controlled for age and education on scores that have not undergone normative correction.

## Results

For the following results, please refer to **Figure 1** as a reference for directionality regarding both the hippocampus and cerebellum.

Cognitive data is primarily derived from V1; however, Visit 2 (V2) cognitive data were used for the Virtual Supermarket Task and 4 Mountains test if data from V1 was missing. V2 was scheduled 6 months after V1 to preserve the sample size. To ensure that temporal lag of testing was not significantly impacting our results, follow-up analyses were conducted with all V2 participants excluded. We found no differences in resting functional MRI (rsfMRI) results. The sole difference we found was in behavioral performance in which patients with a family history of dementia performed better than those without a family history on the 4 Mountains Test. Notably, these analyses were matched for age and sex with a different and smaller subset of the sample to exclude V2 participants as compared to the analyses with a larger cross-sectional sub-sample.

### Cognitive Performance, Sex Differences, and family history of dementia

We discovered several robust associations between cognitive performance, familial dementia, and sex across participants.

Our linear regressions with sex as a predictor and cognitive performance as the outcome—controlling for age and education—revealed significant relationships for story memory delayed recall (β = 0.187, Cohen’s d = .005, pFDR = .040) and the Virtual Supermarket Task (β = -0.346, Cohen’s d = .004, pFDR < .001), respectively. On average, females had higher scores on story memory delayed recall than males (**Figure 2a**) and males had higher scores than females on the Virtual Supermarket Task (**Figure 2b**) across all participants, when controlling for age and education level. There were no significant sex differences seen across the remaining tasks (story memory immediate recall, immediate memory index, delayed memory index, visuospatial/constructional index, and 4 Mountains) after FDR correction (pFDR > .05).

**Figure 2.**
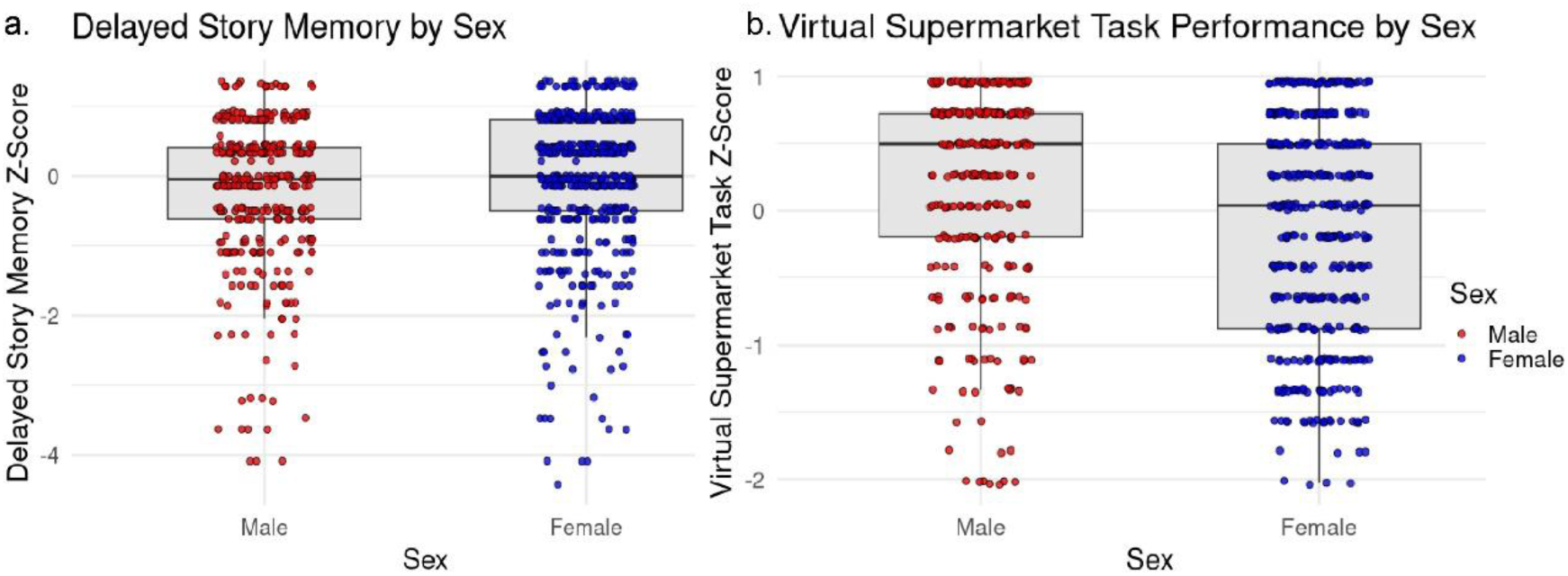
**a.** On average, females had higher scores on story memory delayed recall Z-score than males when controlling for age and education level. **b.** On average, males had higher scores on Z-scored Virtual Supermarket Task than females. These figures depicts raw values and does not depict values adjusted for age and education level.

The unique and combined influences of familial history of dementia and sex on cognitive performance were assessed via interactions (FH group*Sex) and main effects in an age and sex matched sample of FH+ and FH-participants (see Methods section 1.1) in separate linear regressions. We did not find significant interactions of familial history by sex across cognitive tasks (pFDR > .05). However, our regressions did reveal significant main effects for familial history and story memory immediate recall (β = 0.346, Cohen’s d = .012, pFDR = .028; **Figure 3a**) and delayed recall (β = 0.410, Cohen’s d = .013, pFDR = .028; **Figure 3b**) when controlling for age and education level, as well as the immediate memory index (β = 5.159, Cohen’s d = .186, pFDR = .029; **Figure 3c**). Across these results, FH+ participants scored higher than FH-participants. Remaining cognitive measures: coding delayed memory index visuospatial/constructional index, 4 Mountains, and Virtual Supermarket Task did not show main effects of parent history when controlling for age and education level in tasks without normative data correction (pFDR > .05). We are not reporting main effects of sex on cognitive performance as those results have been reported with the full sample as compared to this smaller, age and sex matched sample.

**Figure 3.**
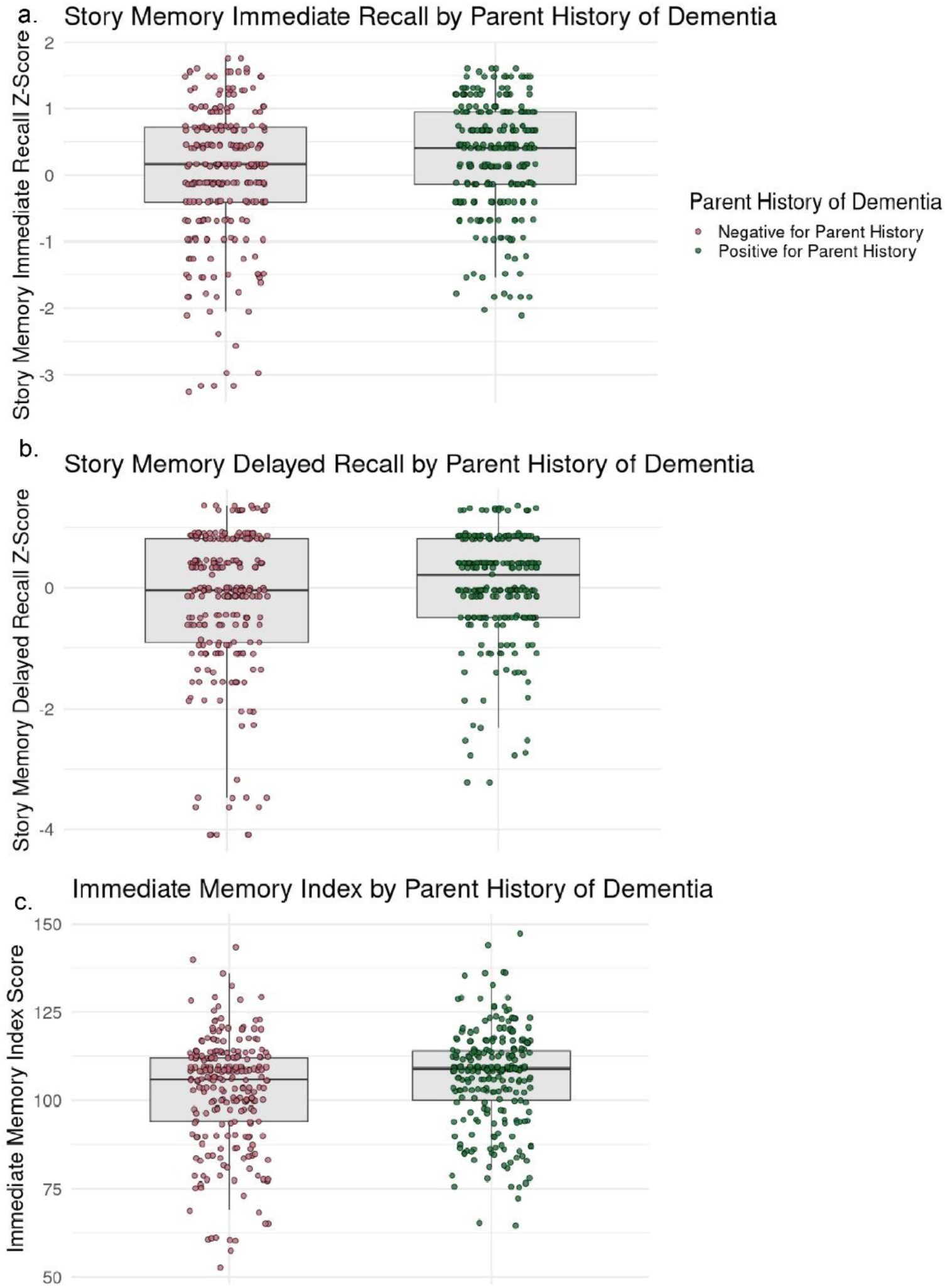
Group differences between FH+/FH-participants. **a.** FH+ participants positive scored higher on story memory immediate recall average than FH*-* participants. **b.** FH+ participants performed better on story memory delayed recall than FH-participants. **c.** FH+ participants performed better on the immediate memory index than FH-participants. Mean and interquartile range are indicated by black lines on each boxplot.

### Cognitive Performance by RBANS Group Differences

Separate linear regressions were conducted to examine the main and interactive effects of RBANS group and sex (RBANS Group*Sex) on cognitive performance. We did not find significant interactions of RBANS Group (i.e., MCI risk versus CN) by sex across cognitive tasks (pFDR > .05). Notably, there were significant main effects for RBANS Group on the 4 Mountains (β = -0.359, Cohen’s d = .370, pFDR < .001; **Figure 4a**) and the Virtual Supermarket Tasks (β = -0.494, Cohen’s d = 1.397, pFDR < .001; **Figure 4b**) when controlling for age and education level. For both tasks, CN participants displayed higher scores, on average, than MCI Risk participants. The remaining cognitive measures, story memory immediate recall story memory delayed recall coding, immediate memory index, delayed memory index, and visuospatial/constructional index, did not show main effects of RBANS group when controlling for age and education level (pFDR > .05). We are not reporting main effects of sex on cognitive performance as those results have been reported with the full sample.

**Figure 4.**
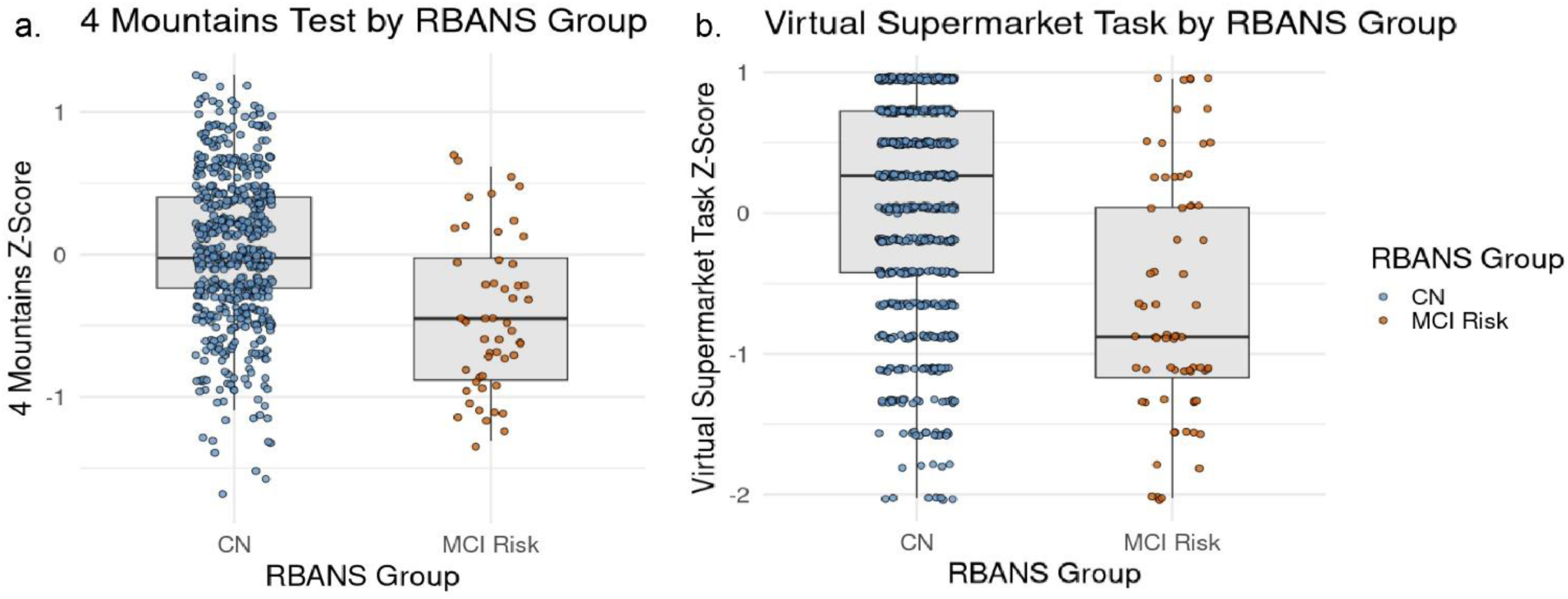
Group differences between participants CN and MCI Risk participants. CN participants scored higher on the 4 Mountains Test than MCI Risk participants when controlling for age and education level. CN participants scored higher on the Virtual Supermarket Task than MCI Risk participants when controlling for age and education level. Raw values uncorrected for age and education level are depicted here.

### Cognitive Performance and CB-HP Connectivity in Familial Dementia History

Complex relationships between CB-HP connectivity, familial dementia, and cognitive performance were explored via interactions (FH*Cognitive performance) and CB-HP in linear regressions. FC data were extracted from CONN toolbox and harmonized to account for multiple scanner site variance using ComBat^74^. Regressions with the harmonized ROI-to-ROI values were conducted in R. FH+ participants were matched by age and sex to FH-participants (see Methods section 1.1). We found that linear regressions between CB-HP FC and the interaction between FH group and cognitive performance (i.e., story memory immediate recall, story memory delayed recall, coding, immediate memory index, delayed memory index, visuospatial/constructional index, 4 Mountains, and Virtual Supermarket Task) were not significant in all ROIs, a subset of ROIs, or when controlling for education in variables without normative correction (pFDR > .05). Further, there was not a main effect of FH group (i.e., no significant differences between FH groups) in our results (pFDR > .05).

### Cognitive Performance and CB-HP Connectivity by Sex

Our analysis did not reveal sex differences in in CB-HP FC (pFDR > .05). Linear regressions between CB-HP FC and sex*cognitive performance were not significant across cognitive tasks when controlling for age and education (pFDR > .05).

When replicating these analyses in our subset of ROIs, we found a significant interaction between sex and the visuospatial/constructional index with Hipp_L_Ant_2 (anterior hippocampus) to R_Lateral_boundary_Xaxis (Crus I) (β = -.002, Cohen’s d = - .260, pFDR = .034; **Figure 5**) in which males displayed a positive relationship between CB-HP FC and visuospatial/constructional index score (i.e., greater connectivity was associated with a higher score). Females displayed a slight negative relationship. No other relationships were significant in this subset analysis (pFDR > .05).

**Figure 5.**
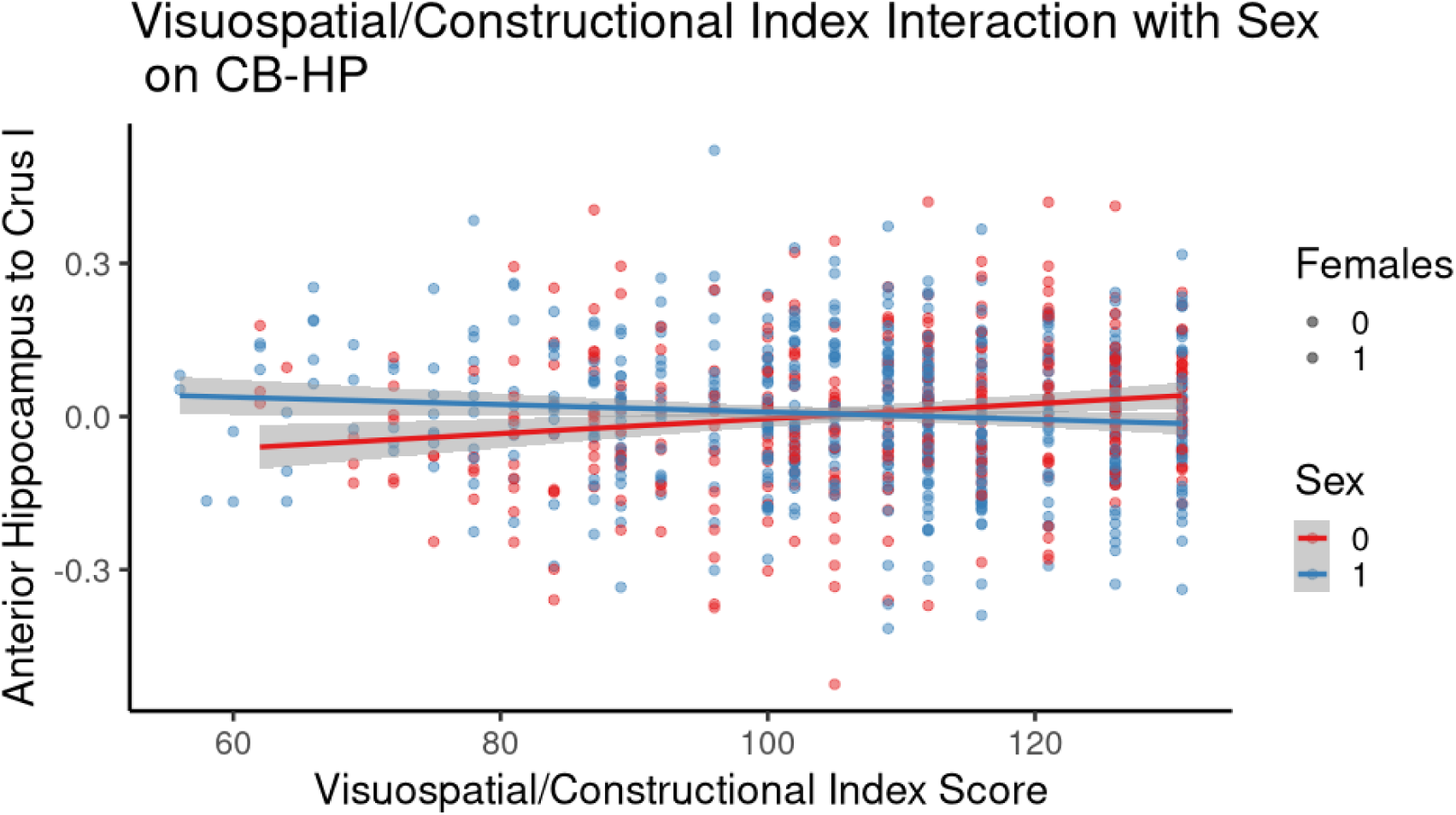
Interaction between Sex and the visuospatial/constructional index with the anterior hippocampus (Hipp_L_Ant_2) to Crus I (R_Lateral_boundary_Xaxis) in which males (red) displayed a positive relationship between CB-HP FC and visuospatial/constructional index score (i.e., greater connectivity was associated with a higher score); whereas, females (blue) displayed a slight negative relationship where lower CB-HP FC was associated with a higher score in all ROIs. The gray indicates a 95% confidence interval.

### Cognitive Performance and CB-HP Connectivity in RBANS MCI Risk and CN Groups

We conducted additional exploratory analyses to assess preclinical cognitive decline using the RBANS total index scores. Established cutoff scores were employed to split groups into MCI risk (<85) and CN (85+)^56^. There were notable differences in group sizes (CN = 748, MCI risk = 104, **Table 1**). There were not significant differences in CB-HP FC between groups (pFDR > .05) across all ROIs or in the exploratory subset of ROIs.

We tested CB-HP connectivity by regressing it on Cognitive Performance, RBANS group (MCI risk vs. CN), and their interaction. There were no significant interactions between RBANS group and cognitive performance with CB-HP FC (pFDR > .05) across all ROIs or in the exploratory subset of ROIs when controlling for age and education level. However, there were significant intracerebellar interactions with behavior (**Supplementary Table 3**). There was a significant interaction for immediate memory, such that CN displayed greater intracerebellar FC with higher immediate and delayed memory indices, whereas MCI risk showed lower FC with higher scores on these tasks (**Supplementary Table 3; Supplementary Figure 2**).

When investigating these interactions in our subset of ROIs, we revealed a significant interaction for RBANS group by story memory immediate recall and Hipp_L_Post2 (posterior hippocampus) to R_CB32 (Crus I) in which MCI Risk participants displayed a positive relationship between CB-HP FC and story memory immediate recall performance and CN participants presented with a slight negative relationship between CB-HP and story memory immediate recall performance (β = .062, Cohen’s d = .247, pFDR = .024; **Figure 6**). There were no additional significant interactions in the subset of ROIs (pFDR > .05).

**Figure 6.**
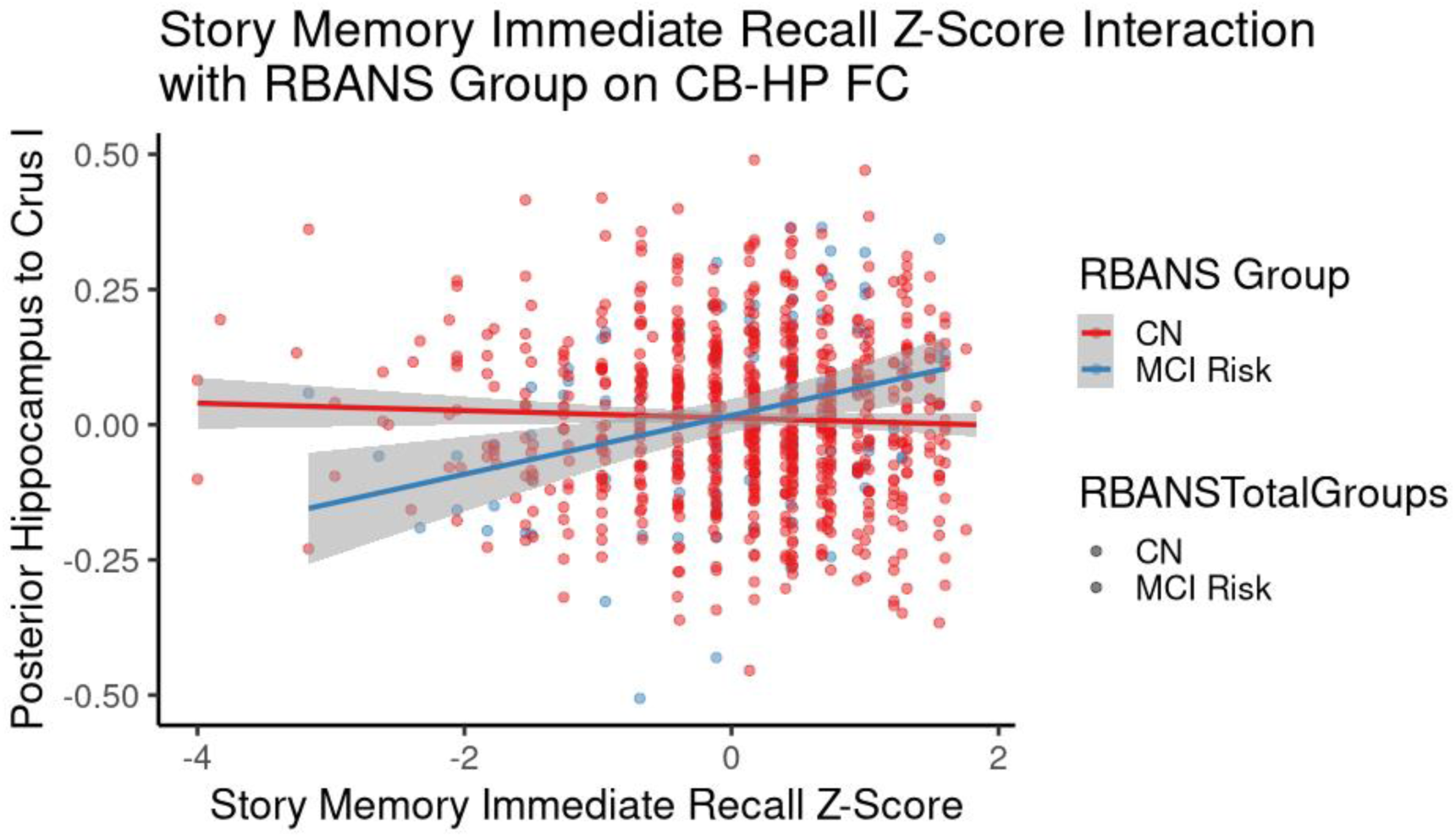
Interaction between RBANS Group and the story memory immediate recall performance with the posterior hippocampus (Hipp_L_Post2) to Crus I (R_CB32) in which CN (red) displayed a negative relationship between CB-HP FC and the story memory immediate recall score (i.e., greater connectivity was associated with a lower score); whereas, MCI risk (blue) displayed a positive relationship where higher CB-HP FC was associated with a higher score in a subset of ROIs. The gray indicates a 95% confidence interval.

### Age, Education Level, and CB-HP Connectivity

Neither age, nor education level were significantly associated with CB-HP connectivity in their respective linear regressions (pFDR > .05).

### Cognitive Performance and CB-HP Connectivity Across Participants

Regarding relationships across all participants between cognitive measures (i.e., story memory immediate recall, story memory delayed recall, coding, Immediate memory index, delayed memory index, and visuospatial/constructional index) and CB-HP FC (all ROIs), the immediate memory index was significantly associated with Hipp_L_Ant_2 (anterior hippocampus) to R_CB25 (Crus I) (β = .001, Cohen’s d = .284 pFDR = .024; **Figure 2**) and the intracerebellar connection between R_CB9 (Lobule VIII) and R_CB18 (Lobule VIII) (β = .002, Cohen’s d = .288, pFDR = .024; **Supplementary Figure 3**). Greater anterior hippocampus to Crus I connectivity was linked to higher immediate memory index scores (i.e., better performance). We did not reveal significant relationships between the remaining cognitive measures and CB-HP FC (all ROIs) when controlling for age and education level (pFDR > .05).

Our exploratory subset (Crus I, Crus II, and hippocampus) of ROI regressions with cognitive performance across participants revealed significant associations between Hipp_L_Ant_2 (anterior hippocampus) to R_CB25 (Crus I) and both the immediate memory index (maintained significant finding; **Figure 7a**) and story memory immediate recall (β = .021, Cohen’s d = .257, pFDR = .038; **Figure 7b**), when controlling for age and education. Better performance on these measures were associated with greater CB-HP FC. Conversely, greater CB-HP was linked to lower Delayed Memory Index scores; Delayed Memory Index and Hipp_L_Ant_1 (anterior hippocampus) to R_CB32 (Crus I) (β = -.001, Cohen’s d = -.263, pFDR = .028; **Figure 7c**). There were no other significant relationships between FC and cognitive measures in the subset of ROIs examined (pFDR > .05).

**Figure 7.**
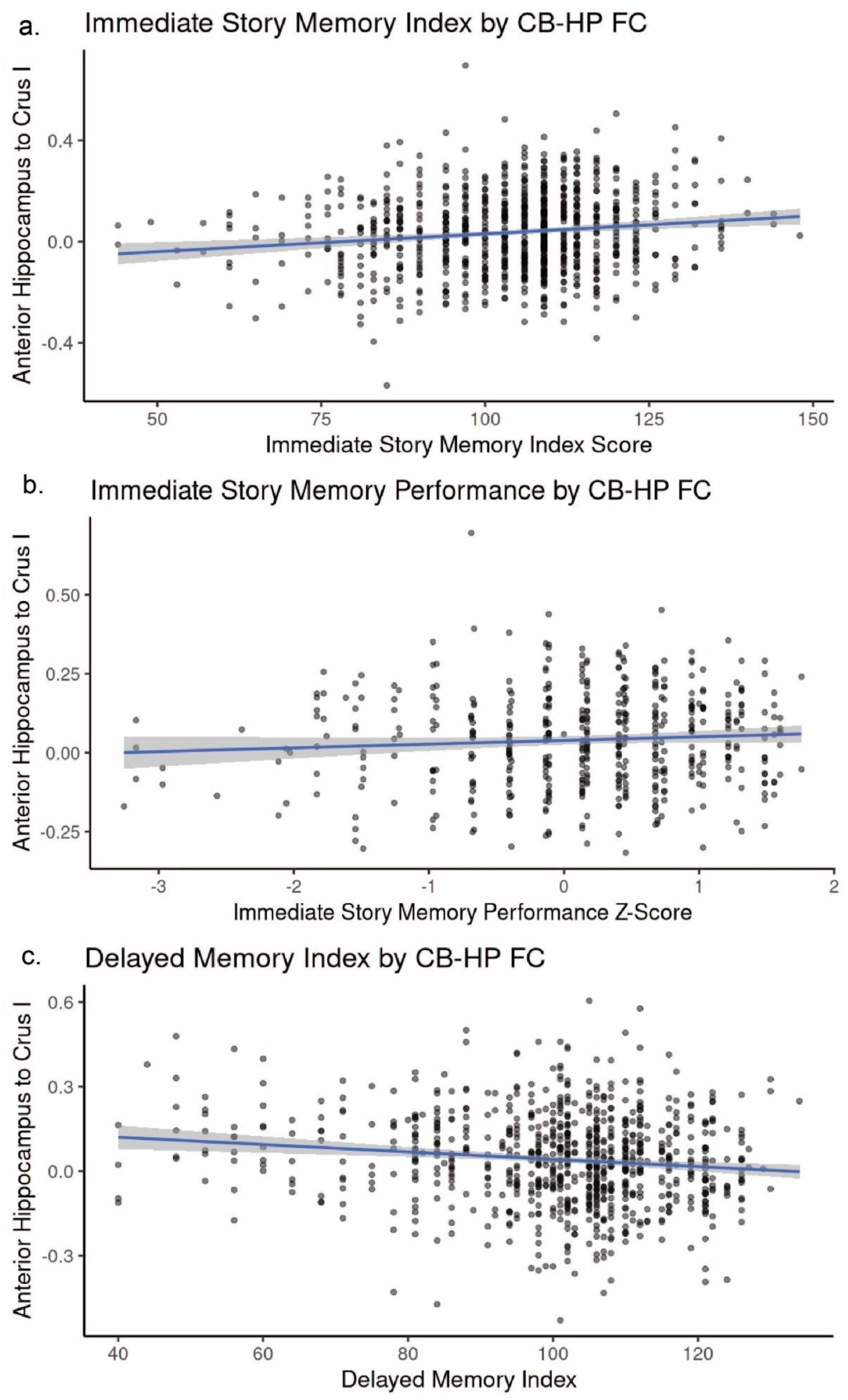
**a.** Greater FC between the anterior hippocampus (Hipp_L_Ant_2) and Crus I (R_CB25) was associated with better immediate recall across several tasks in all participants. **b.** Greater FC between the anterior hippocampus (Hipp_L_Ant_2) and Crus I (R_CB25) was linked to higher z-scored Immediate Story Memory in all participants. **c.** Greater FC between the anterior hippocampus (Hipp_L_Ant_1) to Crus I (R_CB32) showed a negative relationship with delayed memory index scores across all participants in a subset of ROIs. The gray indicates a 95% confidence interval.

## Discussion

Here, we examined relationships between the CB-HP circuit, cognitive performance, and dementia risk in large sample of healthy older adults. Regarding cognitive performance, we found females outperformed males on a delayed episodic memory task, and males outperformed females on a spatial memory task. FH+ participants displayed better immediate and delayed recall for a story as well as short recall for several memory tasks as compared FH-. Those at risk for MCI based on the RBANS displayed poorer performance on both spatial memory tasks. Concerning FC, there were interaction effects such that the relationships between CB-HP FC and behavior differ in males and females. Similarly, MCI risk as defined by the RBANS also had differing relationships between CB-HP FC and behavior. Across this cohort, better immediate recall on memory tasks was associated with greater FC between the anterior hippocampus and Crus I. Conversely, we found that better performance on delayed recall memory tasks was linked to lower CB-HP FC.

This study expands on findings from our previous work showing age and hormone effects as modulators of the cerebello-hippocampal circuit^19^. Further, it adds depth to Paitel and colleagues’ recent study which found lower resting-state FC between whole-region hippocampus to cerebellum with increased age in participants with genetic dementia risk, and higher abnormal AD biomarkers in middle-aged and older adults^18^. They showed greater FC between the hippocampus and cerebellum was linked to better episodic memory performance across their aging sample^18^. Our study offers novel perspectives into the CB-HP circuit and several types of dementia risk factors (i.e., sex, family history of dementia, and RBANS TIS) and adds to the literature linking dementia risk factors to cognitive performance.

### Cognitive Performance, Sex Differences, and family history of dementia

Broadly, the literature has shown patterns of females outperforming males in verbal abilities and males outperforming females in spatial abilities ^78–85^. The current pattern, better delayed story recall in females and better virtual-navigation performance in males, aligns with the longstanding profile of sex differences that remains detectable even in later life. This advantage may provide females with a cognitive “reserve,” delaying a clinical expression of neuropathology despite equivalent amyloid burden ^85^. Specifically, Sundermann et al. found that females outperformed males on verbal memory when amyloid burden was low, and performance was equivalent to males when amyloid burden was high. However, we expected our sex difference findings to extend across our verbal episodic memory (story memory immediate *and* delayed recall) and spatial tasks (coding, visuospatial/constructional index, 4 Mountains, *and* Virtual Supermarket Task). Males have shown advantages with spatial performance in virtual environments^86^ which suggests that the type of spatial task may be driving the sex differences found in our cohort. Our results therefore extend the extant literature by demonstrating that these sex-linked strengths generalize to cognitive tasks in healthy elders, underscoring the need to account for both task domain and participant sex when interpreting cognitive outcomes in aging research.

Contrary to the risk-deficit pattern often found in first-degree relatives of persons with Alzheimer’s disease, our healthy FH+ older adults exhibited superior verbal episodic memory (both for short and longer delays) and immediate recall (verbal and visual) than their FH-peers. Other large cross-sectional studies have reported lower baseline episodic-memory performance among FH+ individuals as measured in early to mid-adulthood^33,87^. One explanation for our findings is cognitive reserve. Qi and colleagues have shown that mid-life cohorts with genetic AD risk who maintain intellectually stimulating activities and higher occupational complexity preserve, or even enhance, episodic memory in later life^88^. Story-recall measures also rely on using the story’s meaning and context to organize what you hear and remember its main ideas, processes that may remain robust until more advanced disease stages. Our findings therefore suggest that parental history of dementia alone is not a consistent predictor of cognitive performance.

### Cognitive Performance by RBANS Group Differences

Groups classified with RBANS Total Index cut-offs (MCI risk and CN) established by Karantzoulis and colleagues^56^ demonstrated differences in cognitive performance on two spatial memory tasks. Namely, CN outperformed the MCI risk group on both the 4 Mountains Test and the Virtual Supermarket Task. These findings align with growing evidence that hippocampal-dependent spatial memory and navigation are especially sensitive to prodromal change^62,63,89–91^. Allocentric place-memory on the 4 Mountains Test distinguishes cross-sectional MCI from CN and also predicts two-year conversion to Alzheimer’s dementia with 93% accuracy^91^. Similarly, Virtual Supermarket Task performance yields excellent discriminative validity for MCI versus healthy aging^90,92^. These domain-specific deficits emerge against a background of otherwise subtle neuropsychological change and appear earlier than traditional word-list measures, a pattern highlighted in a recent review that discusses spatial navigation as an understudied cognitive marker of preclinical Alzheimer pathology^89^. Taken together, our results and past work suggest that RBANS-defined MCI risk captures meaningful variance in hippocampal function that is expressed most clearly when tasks demand allocentric mapping, strategic route planning, and rapid updating of landmark-goal relationships. Embedding such ecologically valid spatial tests within conventional batteries could therefore sharpen risk stratification long before global cognitive scores fall below diagnostic thresholds.

### Cognitive Performance and CB-HP Connectivity in Familial Dementia History

Contrary to our hypothesis, neither family-history status nor its interaction with cognitive performance was linked to CB-HP FC in our cohort. Conversely, cognitively normal offspring of late-onset Alzheimer’s disease (AD) patients show weaker posterior-cingulate to medial-temporal rsfMRI FC^45^. Further, lower cerebrospinal-fluid levels of Aβ₄₂ (a biomarker for AD) in middle-aged FH+ Black Americans is linked to reduced DMN connectivity^44^. Crucially, recent evidence shows that genetic and biomarker risk for AD can modulate CB-HP FC in cognitively unimpaired adults^18^. Namely, Paitel et al. found that those with lower AD risk showed age-related increases in CB-HP FC, whereas those with greater risk for AD exhibited marginal age-related decreases in CB-HP FC; moreover, greater CB-HP FC was associated with better episodic-memory performance across groups^18^. Together with those findings, our results raise the possibility that familial history alone is insufficient to perturb CB-HP communication; additional pathological load (e.g., amyloid, tau, APOE ɛ4 presence) may be required before the circuit’s FC or cognitive contributions are compromised. Longitudinal work will be necessary to determine whether changes in CB-HP FC mark the transition from an at-risk but cognitively intact state to prodromal dementia.

### Cognitive Performance and CB-HP Connectivity by Sex

Our sex by cognitive performance interaction with CB-HP FC analysis revealed stronger FC between the left anterior hippocampus and right Crus I was linked to higher RBANS visuospatial/constructional index scores in males, whereas females showed a slight inverse trend. This pattern dovetails with evidence that the CB-HP circuit contributes to visuospatial abilities^21,23–25^. It also aligns with findings of Crus I– hippocampal synchrony that is recruited specifically when humans navigate using sequence-based or allocentric strategies ^24^ as the visuospatial/constructional index gauges visuospatial planning and spatial orientation. As we previously noted, males tend to outperform females on both standardized visuospatial batteries and virtual navigation tasks. Relatedly, neuroimaging work shows that males rely more heavily on hippocampal circuitry during way-finding, whereas females preferentially engage parietal and prefrontal regions^93^. Together, these findings provide a coherent framework for our interaction: greater male, but not female, visuospatial performance appears to benefit from greater anterior-hippocampus–Crus I FC, consistent with the idea that males capitalize on CB-HP FC for spatial sequencing, whereas females may solve the same tasks via alternative cortical networks. Future work that combines task-based fMRI with strategy probes and hormone measurements alongside this index could clarify whether this sex-divergent FC–performance link reflects differential network recruitment, life-long learning strategies, or modulatory effects of gonadal steroids.

### Cognitive Performance and CB-HP Connectivity in RBANS MCI Risk and CN Groups

Our RBANS group by cognitive performance interactions with CB-HP FC across our cerebellar and CB-HP connections suggest that memory-related communication within the posterior cerebellum shifts from an efficiency framework in CN to a compensatory framework in MCI risk individuals. For the Immediate Memory Index, greater Lobule VI to Crus I and Crus I to Lobule IX connections were associated with better scores in the CN group, whereas MCI risk participants showed the opposite slope. A similar pattern emerged for the Lobule VI and Crus I with the delayed memory index, reinforcing the notion that efficient within-cerebellar synchrony benefits intact networks but becomes maladaptive once prodromal pathology is present. Experimental work has shown that these same lobules boost visuospatial encoding and sequence prediction when task demands are moderate^23,25^. By contrast, MCI risk individuals showed the opposite slope, which is inconsistent with reports that posterior-cerebellar “hyper-connectivity” marks compensatory recruitment as pathology accumulates^94^. Interestingly, posterior hippocampus to Crus I FC was associated with story memory immediate recall, where greater CB-HP FC benefited the MCI risk group yet showed a slight negative relationship in CN. This pattern parallels the way hippocampal-Crus I coherence rises only when rodents engage in demanding sequence-based navigation but not in effortless place-based shortcuts ^21,24^. Together these findings suggest that efficient posterior-cerebellar connectivity optimizes memory in healthy aging, while expected risk for neurodegeneration pushes the system toward broader, less efficient CB-HP recruitment to uphold immediate and episodic recall.

### Age, Education Level, and CB-HP Connectivity

Neither chronological age nor years of formal education were associated with CB-HP FC in our 50- to 88-year-old cohort. This pattern both contrasts with and helps contextualize recent work. Recently, our lab examined a similarly broad adult sample (35-86 y) and reported a negative age–CB-HP slope^19^. Our results highlighted reduced CB-HP coupling as a possible marker of normative network segregation loss. By comparison, Paitel et al. found that age effects were bidirectional with increasing CB-HP FC in APOE ε4 non-carriers and decreasing FC in ε4 carriers or individuals with abnormal tau/amyloid ratios^18^. Because our study did not stratify by genetic or biomarker risk, opposing trajectories within subgroups may have cancelled out, yielding no net association. Earlier rsfMRI work has shown both age-related hypo and hyper-connectivity between the cerebellum and large-scale cortical networks^14^, suggesting that directionality is more related to individual reserve and/or latent pathology. The absence of an education effect is also noteworthy as higher education has been linked to preserved global network segregation in patients at risk for AD^95^, such reserve mechanisms may influence task-evoked efficiency rather than baseline CB-HP synchrony. Taken together, our null findings imply that age and education exert only indirect or subgroup-specific influences on the CB-HP circuit, reinforcing calls for designs that jointly model lifespan, genetic risk, neuropathology and reserve factors when probing this novel pathway.

### Cognitive Performance and CB-HP Connectivity Across Participants

Across the full sample of healthy older adults, stronger FC between the left anterior hippocampus and right Crus I was associated with better immediate recall, whereas a similar hippocampal–Crus I connection predicted poorer performance on delayed recall. Consistent with our positive relationship CB-HP findings, Paitel and colleagues found greater CB-HP FC with better episodic memory performance across participants^18^. Meta-analytic studies likewise place Crus I within a cortico-cerebellar loop that bolsters working-memory and language processes needed to extract and rehearse story gist^96,97^. In contrast, the finding that greater anterior-hippocampus–Crus I connectivity predicts worse delayed recall aligns with reports that hyperactivation marks an inefficient, age-related reliance on networks once consolidation demands increase. One possibility is that heightened CB-HP coordination facilitates initial encoding but, when sustained or excessive, may interfere with the hippocampal re-organization necessary for reliable memory traces, a pattern also seen when over-recruitment of hippocampal redundancy fails to benefit long-term recall in prodromal aging^98^. Longitudinal designs that separate encoding, early consolidation, and retrieval phases will be vital to determine whether these types of results reflect adaptive compensation or the first sign of network inefficiency preceding cognitive decline.

## Conclusion

Posterior-cerebellar and CB-HP networks may support memory via an efficiency-versus-compensation framework, moderate connectivity benefits recall in healthy aging, whereas similar or greater FC appear maladaptive (or compensatory) in individuals at elevated cognitive risk. Sex further modulates these relationships, with males showing patterns of greater CB-HP synchrony with better visuospatial performance. Longitudinal and task-based studies are needed to pinpoint when adaptive FC gives way to inefficient hyper-connectivity and whether this predicts progression to clinical MCI.

## Acknowledgment

This work was supported by R01AG065010 to J.A.B. This work was further supported by the Texas Virtual Data Library (ViDaL), a high-performance cluster, funded by the Texas A&M University Research Development fund. In this cluster the imaging analyses for the current work were carried out using the resources provided by the Texas A&M High Performance Research Computing organization.

